# Accessory genome contributes to the virulence and resistance of the ocular isolate of *Pseudomonas aeruginosa*: A complete genome analysis

**DOI:** 10.1101/497974

**Authors:** Dinesh Subedi, Gurjeet Singh Kohli, Ajay Kumar Vijay, Mark Willcox, Scott A. Rice

## Abstract

Bacteria can acquire an accessory genome through the horizontal transfer of genetic elements from non-parental lineages. This leads to rapid genetic evolution allowing traits such as antibiotic resistance and virulence to spread through bacterial communities. The study of complete genomes of bacterial strains helps to understand the genomic traits associated with virulence and antibiotic resistance. We aimed to investigate the complete accessory genome of an ocular isolate of *P. aeruginosa*.

We obtained the complete genome of the ocular isolate strain PA34 of *P. aeruginosa* utilising genome sequence reads from Illumina and Oxford Nanopore Technology followed by PCR to close any identified gaps. In-depth genomic analysis was performed using various bioinformatics tools. The phenotypic properties of susceptibility to heavy metals and cytotoxicity were determined to confirm expression of certain traits.

The complete genome of PA34 includes a chromosome of 6.8 Mbp and two plasmids of 95.4 Kbp (pMKPA34-1) and 26.8 Kbp (pMKPA34-2). PA34 had a large accessory genome of 1,213 genes and had 543 unique genes not present in other strains. These exclusive genes encoded features related to metal and antibiotic resistance, phage integrase and transposons. At least 24 GIs were predicated in the complete chromosome, of which two were integrated into novel sites. Eleven GIs carried virulence factors or replaced pathogenic genes. A bacteriophage carried the aminoglycoside resistance gene (*aac(3)-IId*). The two plasmids carried other six antibiotic resistance genes.

The large accessory genome of this ocular isolate plays a large role in shaping its virulence and antibiotic resistance.

**Importance:** *Pseudomonas aeruginosa* is an important human pathogen involving many body sites including eyes. The isolates from different infections have a wide variation in their phenotypic and genotypic characteristics despite they have the same host; human. These variations result from the acquisition of genetic elements (accessory genome) from external sources, which are sometimes associated with virulence and antibiotic resistance. This study investigated acquired genes in an eye isolate by analysis of the complete genome. A large portion of the genome of this strain carried accessory genome and many of which had possibly been acquired from environmental bacteria. Notably, these accessory genomes were associated with heavy metal and antibiotic resistance and pathogenesis. This study highlights the importance of genetic study to understand the pathogenesis process of eye *Pseudomonas aeruginosa*.

## Introduction

*Pseudomonas aeruginosa* is associated with many opportunistic and nosocomial human infections such as pneumonia, septicaemia, corneal ulcers (microbial keratitis) and chronic infections in cystic fibrotic lungs (1, 2). Antibiotic resistance in this bacterium is alarmingly on the rise and *P. aeruginosa* has been included in one of the top three priority pathogens urgently requiring new antimicrobial therapies for treatment by the World Health Organization (3). Resistance to different antimicrobials in *P. aeruginosa* is due to its inherent capacity to oppose the action of antibiotics and its capacity to acquire genetic elements that often carry antibiotic resistance genes (4).

The study of the mechanisms used by different strains of *P. aeruginosa* to become resistant to antibiotics or acquire virulence have benefited from genome sequencing and these have identified a number of mobile genetic elements (MGEs) (5-7). However, many of the genomes investigated are not completely closed and, in this context, accessory genes associated with resistance or virulence in addition to mutations may be missed. This may limit the understanding of how resistance and virulence genes are acquired as well as their prevalence in general (8). For example, less than 5% of *P. aeruginosa* draft genomes are complete; furthermore, only 37 plasmids of the total (3003 as of 15/10/2018) in the NCBI *P. aeruginosa* database are complete. Additionally, many of the isolates sequenced are derived from cystic fibrosis infections, while relatively few ocular isolates are represented. This is an important source of genetic information for *P. aeruginosa* as specific subpopulations are thought to be associated with microbial keratitis (9). Notably, those subpopulations are predominantly characterised by possession of genes associated with type IV pili twitching motility, cytotoxicity (*exoU*), and certain genomic islands (9). Indeed, to date, the complete genome of only one *P. aeruginosa* keratitis isolate has been published (10). Very little information is available about the accessory genomes of ocular isolates.

*P. aeruginosa* strain PA34 (referred to as PA34 hereafter) is a multi-drug resistant microbial keratitis isolate which is resistant to gentamicin, imipenem, ciprofloxacin and moxifloxacin. Analysis of its draft genome revealed that PA34 belongs to sequence type 1284 based on multi-locus sequence typing and carried at least 12 acquired resistance genes and possessed the *exoU* gene (11), an important effector gene in the pathogenesis of microbial keratitis (12). A class I integron (In*1427*) that carries two antibiotic resistance genes has been shown to be integrated into a Tn3-like transposon carried by PA34 (13). However, the genetic context of these resistance and pathogenic genes has not been completely elucidated. Hence, in this study we sought to analyse the complete genome of PA34, to examine the genetic structure of accessory genome and to compare the complete genome of PA34 with other genomes of *P. aeruginosa* from the public database.

## Methods

### Bacterial strain and microbiology

The strain *Pseudomonas aeruginosa* PA34 was isolated in 1997 from the cornea of a microbial keratitis patient in a tertiary eye care centre in India (L.V. Prasad Eye Institute, Hyderabad, India). The cause of microbial keratitis was recorded as trauma induced by a stone in the right eye of a 21-year old male. The strain was obtained from the institutional repository without identifiable patient data, following institutional guidelines. Three complete genomes of *P. aeruginosa* were used for the reference: I) *P. aeruginosa* PAO1, which has the most complete and curated annotations (14, 15), II) *P. aeruginosa* PA14, an *exoU* positive strain (16) and III) *P. aeruginosa* VRFPA04, an Indian microbial keratitis isolate (10).

The minimum inhibitory concentration (MIC) to three heavy metals (mercury, copper and cobalt) was determined by broth dilution methods as described elsewhere with minor modification (17, 18). PA34 and PAO1 (as the control strain) were examined against Hg^++^, Cu^++^ and Co^++^ by using twofold serial dilutions from 16mM – 0.31mM. The metal salt solutions (HgCl_2_, CuCl_2_ and CoCl_2_) were prepared in deionised water, sterilized by membrane filtration and added to Mueller-Hinton broth (Oxoid Ltd, Hampshire, UK) to obtain the required concentrations. The experiments were performed in triplicate and repeated three times.

Cytotoxicity was examined by the trypan blue dye exclusion assay (19). Briefly, human corneal epithelial cells (HCEC) were grown in 96 well plates in the presence of SHEM (DMEM (Gibco, Grand Island, NY, USA) supplemented with 10% fetal bovine serum (Gibco), 1.05mM CaCl_2_, 0.5% DMSO, 2ng/mL epidermal growth factor (Gibco), 1% ITS-X (Gibco)). 200μL of 5x10^5^ CFU/mL *P. aeruginosa* in SHEM was exposed to confluent HCEC. Following 3 h incubation at 37°C, the cells were stained with 0.4% trypan blue. The stained cells were observed by microscopy and photographs were taken for the quantitative determination of cytotoxicity. *P. aeruginosa* PAO1 (a non-cytotoxic strain) were used as a control. The experiment was performed in triplicate and repeated three times.

### MinION sequencing and complete genome assembly

Genomic DNA was extracted using a Wizard^®^ Genomic DNA Purification Kit (Promega, Madison, WI, USA) as per the manufacturer’s protocol. The amount of extracted DNA was determined using a Qubit fluorometer (Life Technologies, Carlsbad, CA, USA). The library was prepared using the Ligation Sequencing Kit 1D R9 Version (Oxford Nanopore Technologies Ltd [ONT], SQK-LSK108), as per the manufacturer’s instructions. The sequencing library was loaded into the Flow Cell Mk 1 Spot-ON (ONT, FLO-MIN 107 R9) of the MinION^™^ system using Library Loading Bead Kit R9 Version (ONT, EXP-LLB001) according to the manufacturer’s instructions. The raw reads were obtained using the MinKNOW v1.7.14 (ONT) in a 24-h run experiment and base calling was performed using Albacore v2.0.2 (ONT). A total of 50,831 reads ranging from 174bp to 58,887bp (average = 3,292bp) were obtained. The sequence reads from Illumina (which was obtained using MiSeq (Illumina, San Diego, CA) generating 300bp paired-end reads (11)) and MinION data were assembled using three hybrid assembly pipe-lines; Unicycler v0.4.3 (20), Japsa v1.8.0 (21) and Spades v3.12.0 (22). Spades resulted in the best assembly in terms of contig numbers (Table 1). The genome coverage was calculated using BBmap v35.82 (23).

**Table 1.**
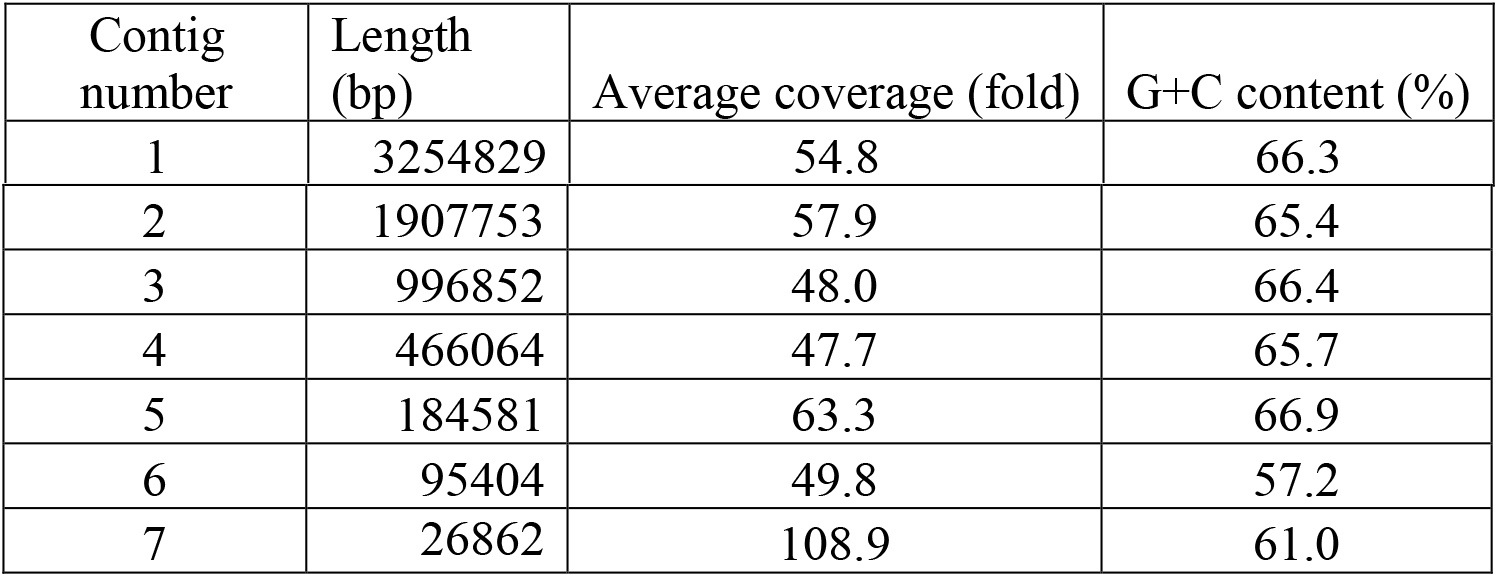
Assembly result from Spades.

All contigs were reordered with reference to the three complete genomes of *P. aeruginosa*. (PAO1, PA14, and VRFPA04) using MUMmer v3.0 (24). Gaps between the contigs were determined by ABACAS v1.3.1 (25) using the “circular reference genomes” flag. Pseudogenomes of different sizes were observed after comparison with different reference strains and in all comparisons, contigs 6 and 7 were observed as unused contigs indicating they are not part of the chromosome. This was further verified by performing a BLAST of unused contigs against reference genomes. The fewest gaps between contigs in the PA34 genome (99 bp) were observed when PA14 was used as the scaffold compared to the other strains. Further, the size of the PA34 pseudogenome was slightly less than the size of the draft genome (~6.8 Mbp) (11) when using PA14 as the scaffold genome. A gap of 99 bp is an indication of a joining point between two contigs (25). Therefore, this comparison was considered as the best and was further used to complete the genome of PA34. Primers were designed for each gap start positions (Supplementary file S1) using Primer3web v4.1.0 (26) and the gap size was verified by PCR. Using this approach, we were able to close the gaps between contigs to generate a single contiguous chromosome based on contigs 1-5. Contigs 6 and 7 had a lower G+C content than rest of the contigs and showed similarity with different plasmids based BLASTn searches against NCBI database. Contigs 6 and 7 were therefore considered to represent two different plasmids and were both complete, circularised elements, referred to here as pMKPA34-1 and pMKPA34-2.

### Bioinformatic analysis

The complete chromosomal genome was annotated using NCBI Prokaryotic Annotation Pipeline (27). Plasmids were annotated by Prokka v1.7 (28) followed by manual examination and curation using information form ISsaga (29), the Rapid Annotations using Subsystems Technology (RAST v2.0) (30) and NCBI BLASTn searches. Genomic islands were predicted on the basis of MAUVE (31) whole genome alignment against *P. aeruginosa* strains PAO1, PA14 and VRFPA04. Through this comparison, DNA blocks of four contiguous open reading frames were predicted as genomic islands of PA34 if these blocks were present in PA34 but absent in anyone of the three reference genomes used for comparison (5). Pangenome analysis was conducted using Roary v3.12.0 (32), and all four genomes (PAO1, PA14, VRFPA04 and PA34) were annotated using Prokka to avoid annotation biases. Other software used for analysis were CRISPRCasFinder database (33) CGview (34), Easyfig (35), SnapGene Viewer v4.3.2 (36), VENNY v2.1.0 (37) and Resfinder V3.0 (38). The nucleotide sequence of the complete chromosome and two plasmids were made available in the NCBI database under GenBank accession numbers CP032552, MH547560 and MH547561.

## Results and Discussion

### 1. General features of *P. aeruginosa* PA34 genome

The statistics of the complete genome of *P. aeruginosa* PA34 (Table 2) were broadly matched with other published complete genomes of *P. aeruginosa* (39). A total of 6462 coding sequences (CDS) were predicted in the 6.8 Mbp genome of PA34 using NCBI Prokaryotic Genome Annotation Pipeline (PGAP) (40). Of the predicted CDS, 6314 were predicted to form functional proteins and 148 were pseudogenes.

**Table 2.** Genomic features of *P. aeruginosa* PA34.

| Characteristics |  |
| --- | --- |
| Genome size in base pairs | 6,810,079 |
| G+C content % | 66.1 |
| Number of genes | 6,544 |
| Number of coding sequences (CDS) | 6,462 |
| Number of protein coding genes | 6,314 |
| Number of pseudogenes | 148 |
| Total RNA genes | 82 |
| complete 5S, 16S, 23S rRNAs | 4, 4, 4 |
| tRNAs | 65 |
| ncRNAs | 5 |

Genomic comparison with three reference genomes (PAO1, PA14 and VRFPA04) and the genome of PA34 estimated there were 7643 orthologs in the pan genome. Of those orthologs, 5078 were common amongst all four strains and are suggested to represent the size of the core genome. PA34 had the largest accessory genome compared to these reference strains (having 1213 genes) that had 543 genes unique to PA34. These exclusive genes were determined to encode features related to metal and antibiotic resistance, phage integrase and transposons. Furthermore, in the context of individual reference genomes, PA34 has 886, 737, 946 genes with no orthologs in PAO1, PA14, and VRFPA04, respectively (Fig 1). PA34 shared 124 orthologs with VRFPA04, an eye isolate and those genes were associated with chloramphenicol resistance, pathogenesis (type IV secretion system) and phages (See supplementary file S2 for full annotations of exclusive orthologs).

**Fig 1.**
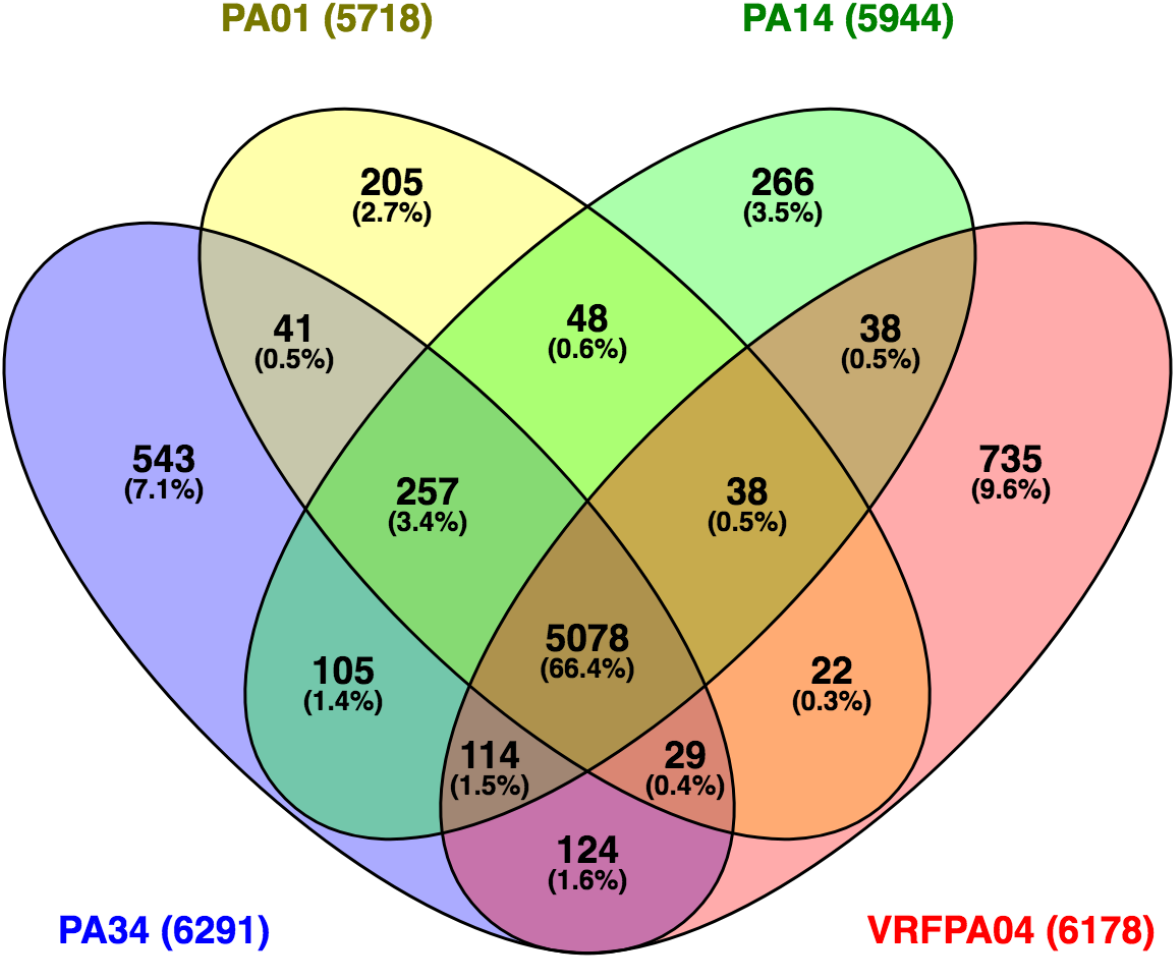
Graphical representation of genomic comparison of *P. aeruginosa* strains. Venn diagram showing the number of common and exclusive orthologs between and amongst four strains of *P. aeruginosa*. The total number of genes examined of each strain is shown in parentheses. The number were predicted by Roary pangenome analysis and figure was created using VENNY v2.1.

No significant CRISPR and associated Cas genes were observed in PA34 based on comparison with the CRISPRCasFinder database (33). CRISPR-Cas system may be negatively correlated with the size of the accessory genome and acquired antibiotic resistance genes because this system restricts the invasion and incorporation of MGEs. Thus, strains without a CRISPR-Cas system are expected to have a larger genome size (41).

### 2. Genomic islands

Genomic islands (GIs) refer to blocks of horizontally acquired DNA that integrate into certain loci in the core genome (42). Such loci are known as Regions of Genomic Plasticity (RGPs). GIs should have a minimum of four contiguous open reading frames, which are not present in any of the set of genomes to which they are compared (5). More than 80 RGPs have been identified in the genome of *P. aeruginosa* (43). The position of RGPs in the genome are indicated by homologous flanking loci in PAO1 (5). RGPs constitute a major portion of the accessory genome and are essential for the adaption of *P. aeruginosa* in diverse habitats. In this study, 24 RGPs were observed in the complete genome of PA34 (Table 3 and Fig 2) when compared with genomes of *P. aeruginosa* strains PAO1, PA14 and VRFPA04. Mathee *et al*. reported 27 to 37 RGPs in individual genomes of five *P. aeruginosa* strains (5). The difference in number of observed RGPs may be due to a difference in the number of genomes used for comparison. Nevertheless, out of 24 GIs in the current study, two GIs were integrated into loci which have not been reported previously (43) and were deemed as GIs new1 and new2 in this study (Table 3). The GI new1 was 68.9 Kbp, was integrated between PA2858 and PA2859 homologues of PAO1 and carried genes for mercury and chromate resistance. The GI new2 had a size of 39.9 Kbp, encodes genes associated with phages and was integrated into loci flanking 4856/4857 of PAO1. In addition, three GIs (RGP23, RGP56 and RGP84) carried phage related genes; phages can be associated with virulence (44). Our analysis showed that at least five GIs were associated with replacement or insertion of pathogenic genes and one GI (RGP23) carried an acquired aminoglycoside resistance gene (*AAC(3)-IId*). Three GIs (RGP2, RGP50, and RGP57) were associated with the type IV secretion system.

**Fig 2.**
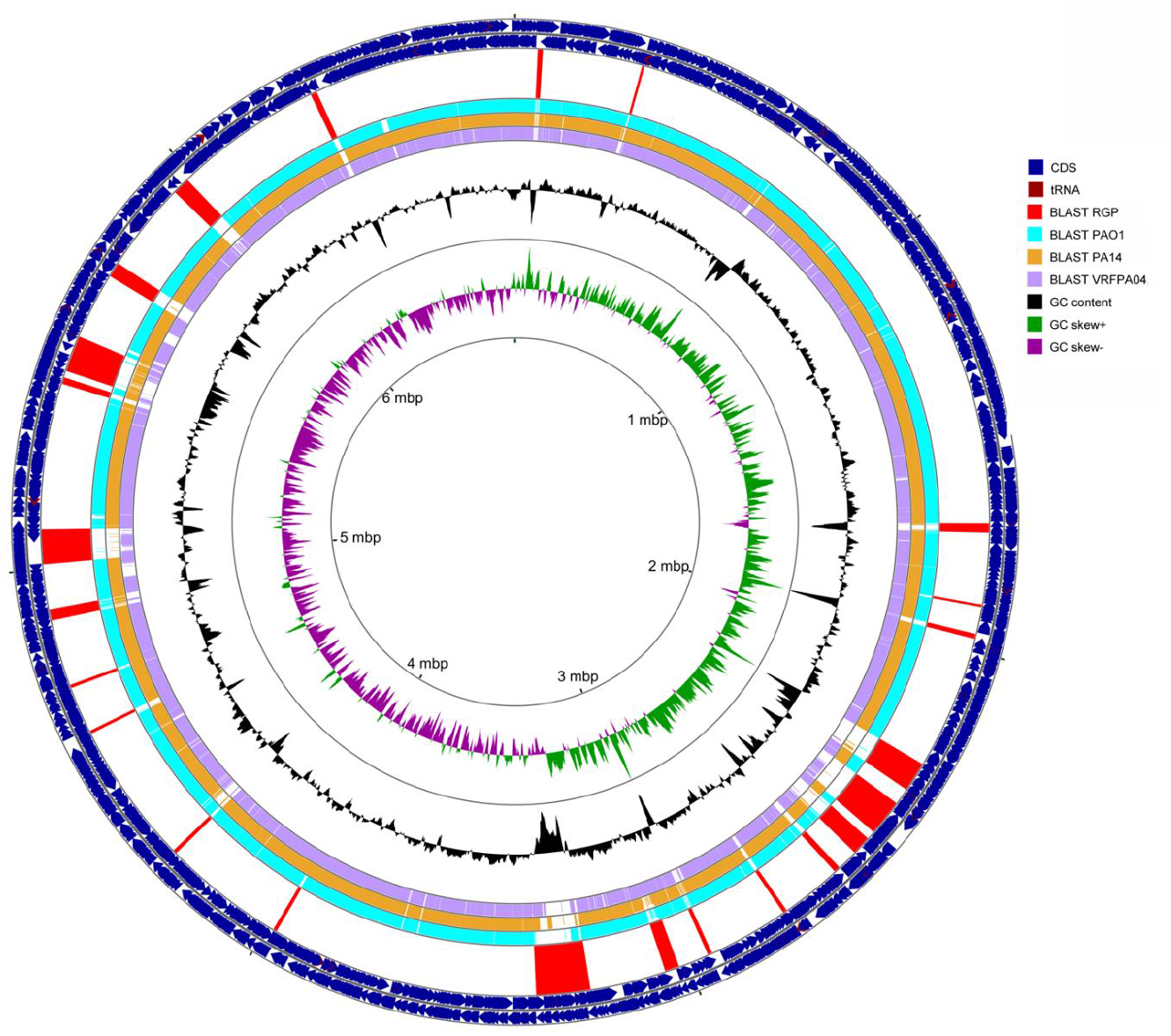
The chromosomal map of *P. aeruginosa* PA34 and position of genomic islands. Circles from outer to inner represent; circle 1. positions of Coding Sequences (CDS) in plus strand (blue), 2. positions of CDS in minus strand (blue), 3. Location of region of Genomic Plasticity (RGPs) (red), in the same order as listed on Table 3, 4. BLASTn comparison against PAO1 (green) 5. BLASTn comparison against PA14 (orange), 6. BLASTn comparison against VRFPA04 (purple). 7. G+C content and deviation from the average, 8. G+C skew in green (+) and purple 9. scale in kbp.

**Table 3.**
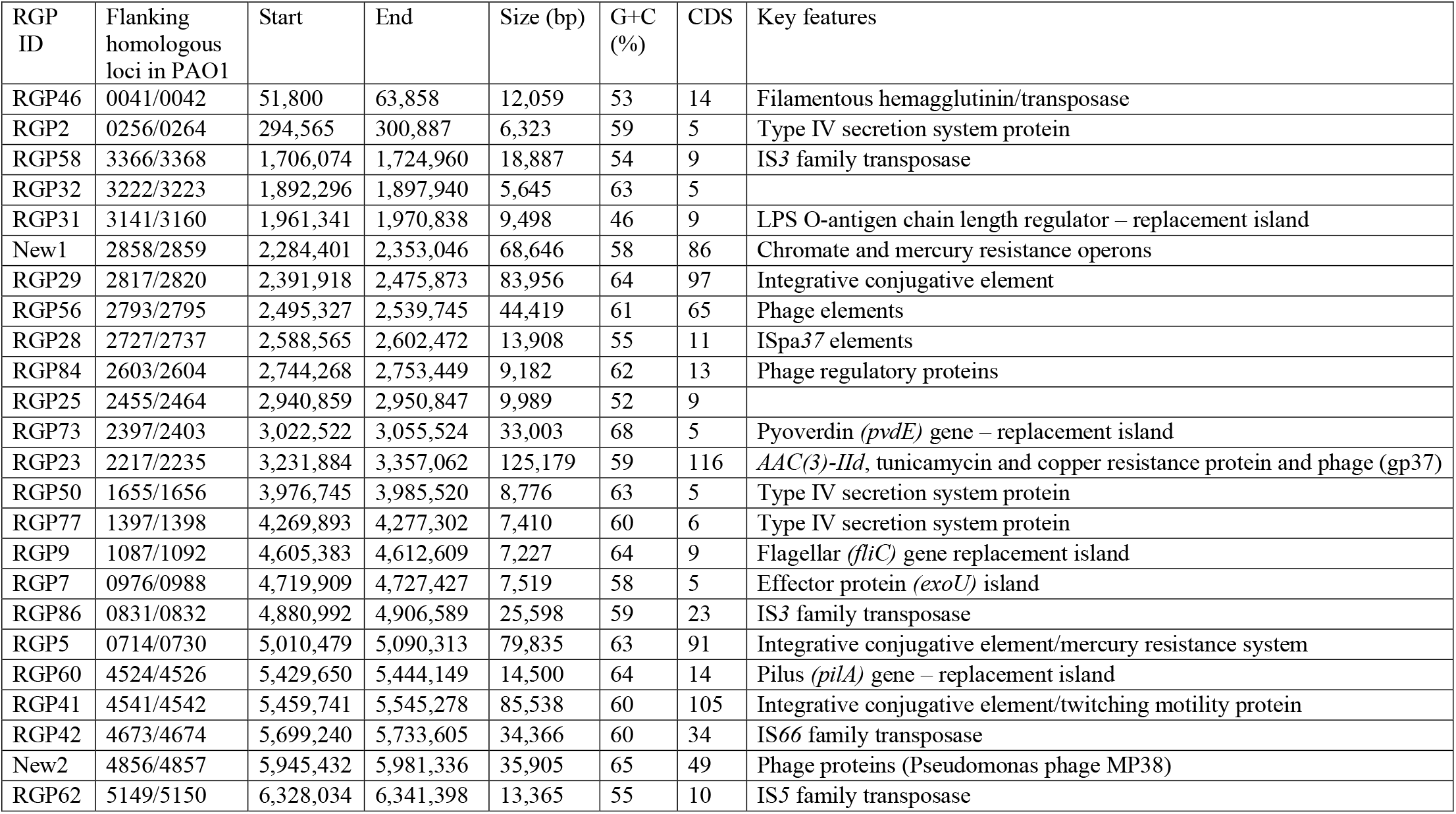
Regions of genomic plasticity (RGP) (5, 43) with associated features in PA34.

Integrative conjugative elements (ICEs) are chromosomally integrated self-transmissible MGEs that can exist as circular extrachromosomal elements and are antecedents of various GIs (42). Our analysis revealed three ICEs in the complete genome of PA34. The ICEs observed in RGP5 and RGP29 were related to *clc-like* ICE elements (Genbank Accession number AJ617740). The parental *clc* element contains genes for chlorocatechol (clc) degradation (45). However, these genes were lost from both RGP5 and RGP29 of PA34. Nevertheless, genes required for integration and conjugation were identified in both of the ICEs in RGP5 and RGP29 (Fig 3A) indicating that these elements may have the capacity to transfer within and between species. This may be the reason for observing *clc*-like GIs in two different loci in this strain. In addition, RGP5 carried mercury resistance genes, which are not present in the parental *clc* elements. This finding suggests that the clc-like elements may incorporate antibiotic resistance genes during genetic recombination.

**Fig 3.**
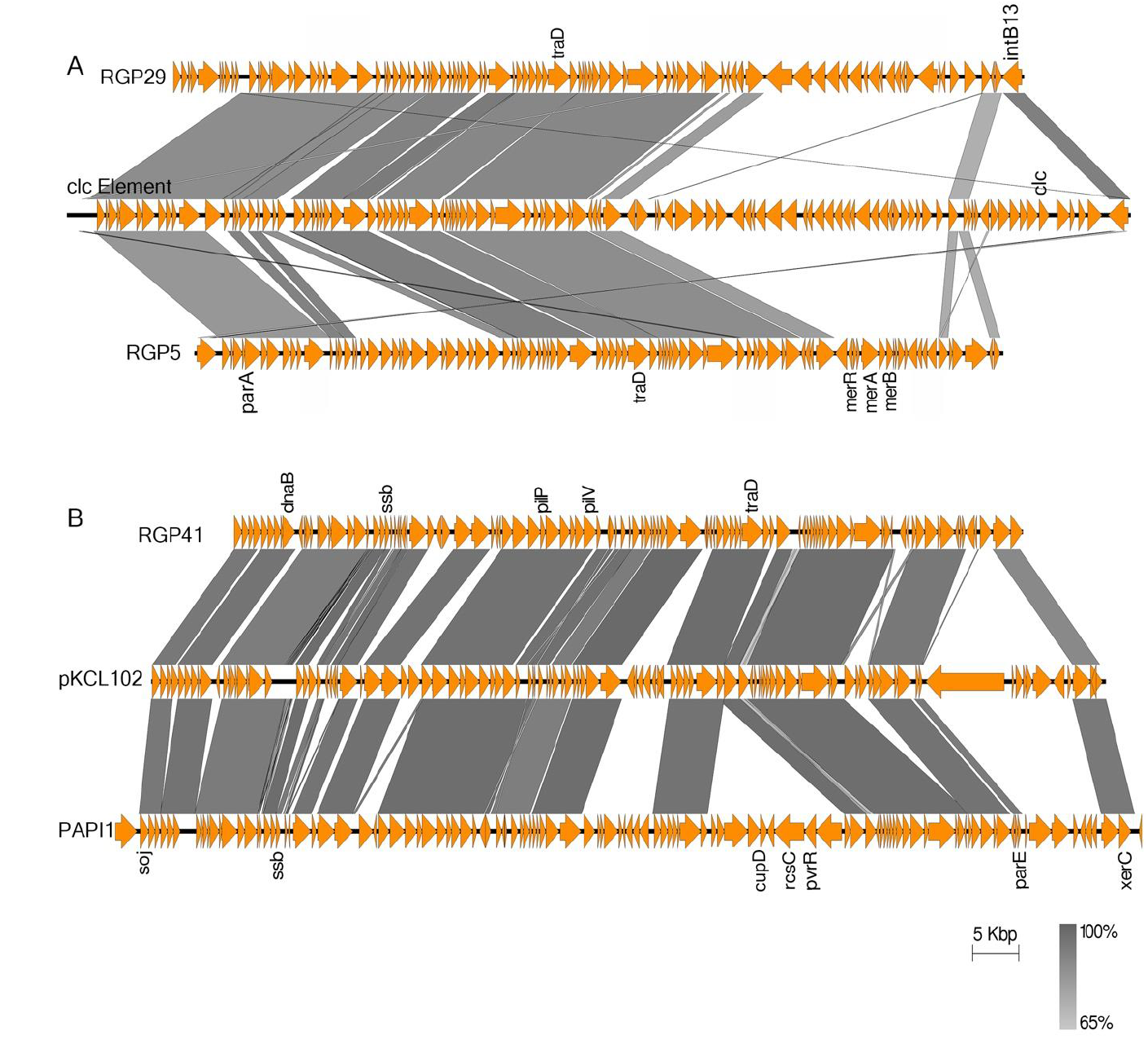
Graphical representation of BLASTn comparison of integrative conjugative elements of PA34. Protein-coding regions are represented by the orange arrows and key features/associated genes are shown. The gradient grey shading represents regions of nucleotide sequence identity (100% to 65%) determined by BLASTn analysis. Figures are drawn to scale using Easyfig (35). (A) A comparison of two genomic islands (RGP 29 and RGP5) of PA34 with a clc-like integrative conjugative element (Genbank Accession number AJ617740) (*parA*= plasmid partition protein, *traD*= conjugative factor, *intB13*=phage integrase, *merA/B/R*= mercury resistance operon, *clc* = chlorocatechol (clc) degradation). (B) A comparison of the genomic island RGP41 of PA34 with pKLC102 (GenBank Accession number AY257538) and PAPI1(GenBank Accession number AY273869) (*soj*= chromosome partitioning protein, *dnaB*= replicative DNA helicase, *ssb*=single strand binding protein, *pilP/pilV*= type IV pilus biogenesis and transfer, *traD*= conjugative factor, *cupD/rcsC/pvrR* = cell surface fimbriae operon, *parE*=plasmid stabilisation protein, *xerC*=integrase)

The third ICE observed in RGP41 of PA34 was related to the pKLC102 family (GenBank Accession number AY257538) and was similar to *P. aeruginosa* pathogenicity island PAPI-1 (GenBank Accession number AY273869) first identified in *P. aeruginosa* clone C (Fig 3B). PAPI-1 is self-transmissible and carries virulence factors such as genes for type IV pilus biogenesis and the *cupD* gene clusters (46, 47). The *cupD* genes are essential for the formation of cell surface fimbriae that are involved in biofilm formation (48). Interestingly, the *cupD* gene cluster appears to have been lost from the RGP41 of PA34, although the genes for type IV pilus biogenesis are still present.

In addition, various members of pKCL102 ICE family are found to be associated with the carriage of *exoU/spcU* genes (42). These GIs have frequently been referred as *exoU* islands such as exoU island A, exoU island B, exoU island C and PAPI-2 (46, 49). Possession of the *exoU* gene, that encodes the type III secretion system effector cytotoxin ExoU, markedly enhances the virulence of *P. aeruginosa* (50). An *exoU*-island of 7.5 Kbp carrying five open reading frames was observed in PA34 (RGP7). In contrast to other *exoU*-islands that contain genes associated with integration, transposition, conjugations and sometimes antibiotic resistance (51), the *exoU* island observed here contains only three genes that have homology to a site-specific integrase (Fig. 4A). The presence of both PAPI-1 and PAPI-2 like elements that act synergistically in virulence (52) may indicate an enhanced virulence of PA34.

**Fig 4.**
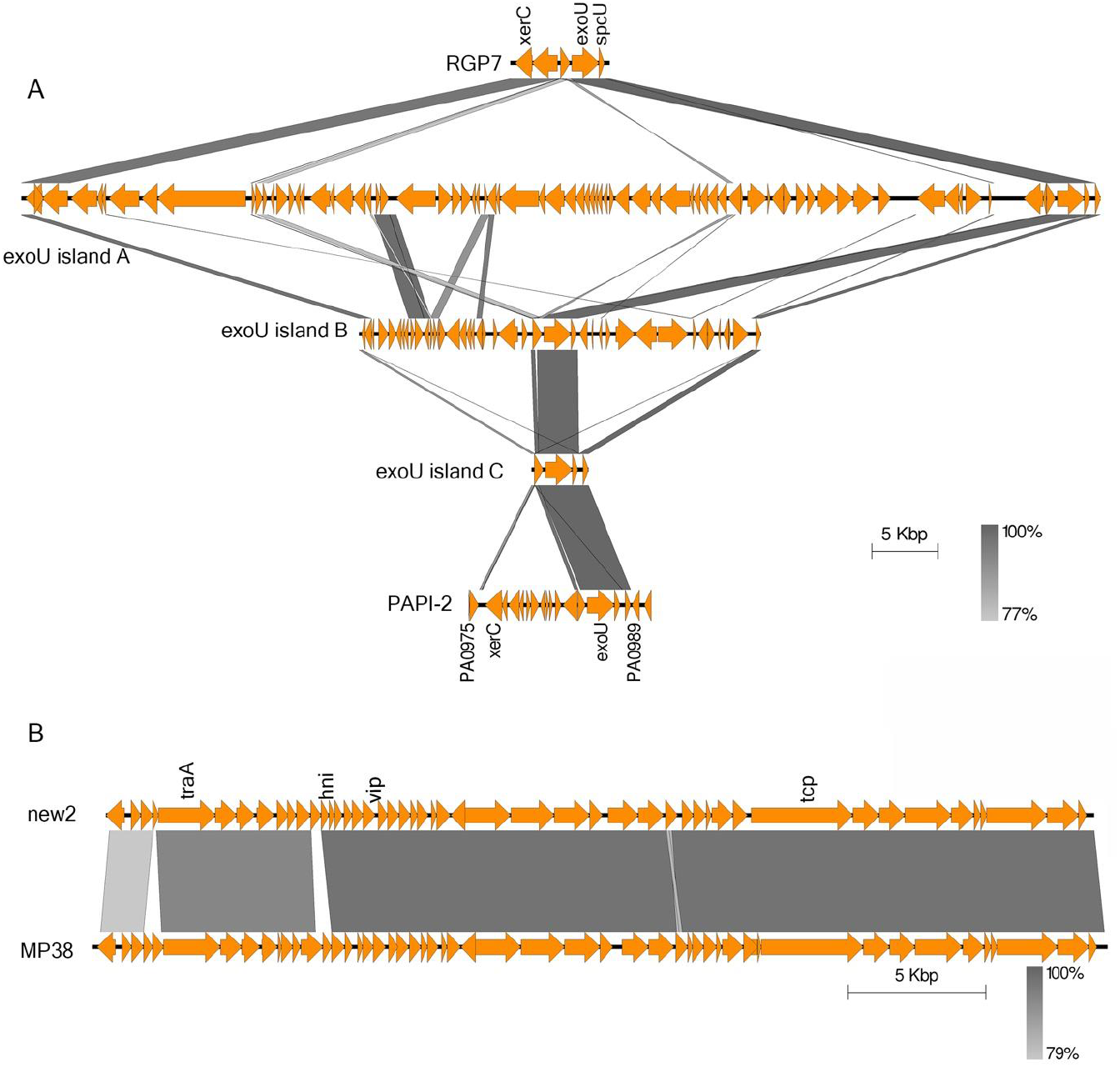
Graphical representation of BLASTn comparison of various genomic islands of PA34. Protein-coding regions are represented by the orange arrows and key features/associated genes are shown. The gradient grey shading represents regions of nucleotide sequence identity (100% to 79%) determined by BLASTn analysis. Figures are drawn to scale using Easyfig (35). (A) A comparison of exoU-island (RGP7) of PA34 with four exoU-islands of different strains (exoU island A, exoU island B, exoU island C and PAPI2) (*xerC*= site specific integrase, *exoU*= type III secretion system effector protein ExoU, spcU= ExoU chaperon protein, PA0975/PA0989=franking loci in PAO1) (B) A comparison of a genomic island (new2) of PA34 with Pseudomonas phage MP38 (GenBank Accession number NC_011611) (*traA*= transposase A, *hni*= host nuclease inhibitor, *vip*= virion protein, *tcp*=tail component protein).

To determine whether the *exoU* gene was functional, cytotoxicity of PA34 was examined in a human corneal epithelium cell line and compared with *P. aeruginosa* PAO1 which is a non-cytotoxic invasive strain (53). Microscopic examination after staining with trypan blue indicated that PA34 was highly cytotoxic (Fig 5) showing a large number of dead cells. Based on the percentage of dead cells, maximum cytotoxicity (of score 4) was observed in PA34 (19). This finding suggests that PA34 *exo U* in RGP7 was transcribed and translated into a functional protein that retained cytotoxicity. The finding also demonstrates that the size of the exoU island is not be associated with activity.

**Fig 5.**
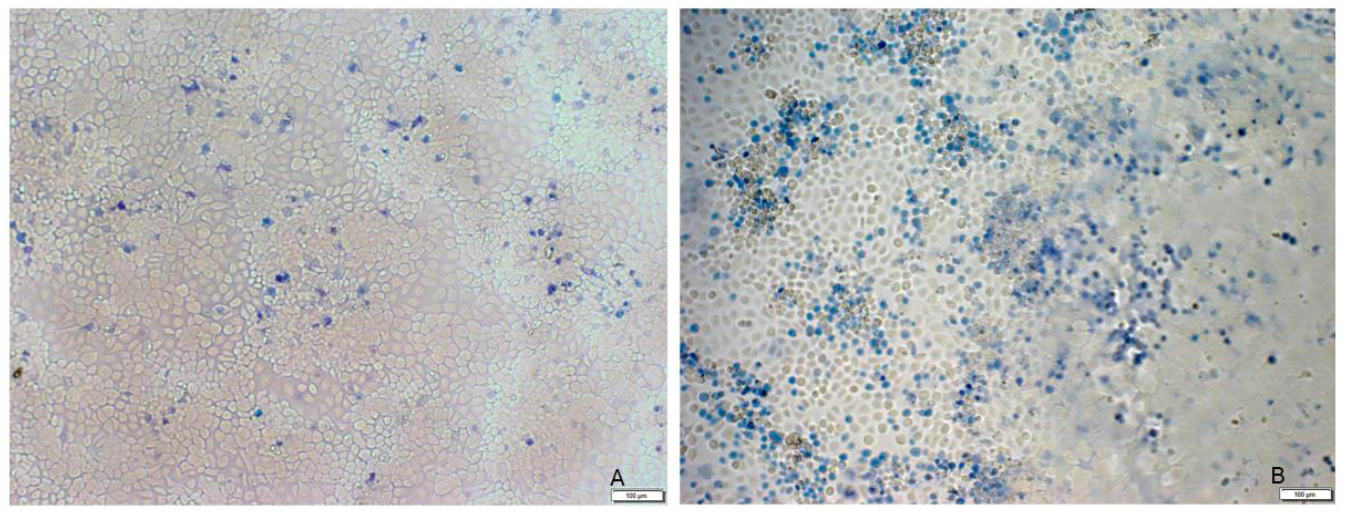
**Microscopic photograph of cytotoxic effect** of (A) *Pseudomonas aeruginosa* PAO1 (the control strain) and (B) *Pseudomonas aeruginosa* PA34 for human corneal epithelium cells strained by trypan blue. Dead cells are strained with trypan blue.

A mercury resistance operon was also observed in the new1 RGP that also carried chromate resistance genes and features related to transposition and conjugation. A BLASTn search against the plasmid database in NCBI did not show significant sequence similarity with new1. However, 51% of the new1 sequence was loosely identical (205 matches with an average identity of 97%) to a the complete genome of *Pseudomonas stutzeri* strain 273 (GenBank accession number CP015641.1). *Pseudomonas stutzeri* is an environmental organism and has the capacity to degrade pollutants such as toxic metals (54). These observations indicate that the new1 may have been acquired by horizontal transfer from organisms in mercury polluted environments. The ability of new1 to transfer between strains needs to be confirmed by further experiments that will also help to confirm the new1 RGP loci is mobile.

In order to compare the phenotypic susceptibility to heavy metals, MIC to Hg^++^, Cu^++^ and Co^++^ were examined. High mercury tolerance was observed in PA34 in comparison to PAO1 (Fig 6), possibly due to the presence of mercury resistance operon in two different GIs (RGP5 and new1) in PA34. Copper tolerance was slightly higher in PA34 than PAO1. Although bacteria require copper as a cofactor for many metabolic processes and can tolerate low concentration of Cu^++^, decreasing sensitivity to copper can be associated with acquisition of copper resistance genes (55). The presence of a copper resistance operon in RGP23 in PA34 may be associated with higher MIC to Cu^++^. On the other hand, cobalt tolerance, for which no acquired genes were observed in this study was low in both PA34 and PAO1.

**Fig 6.**
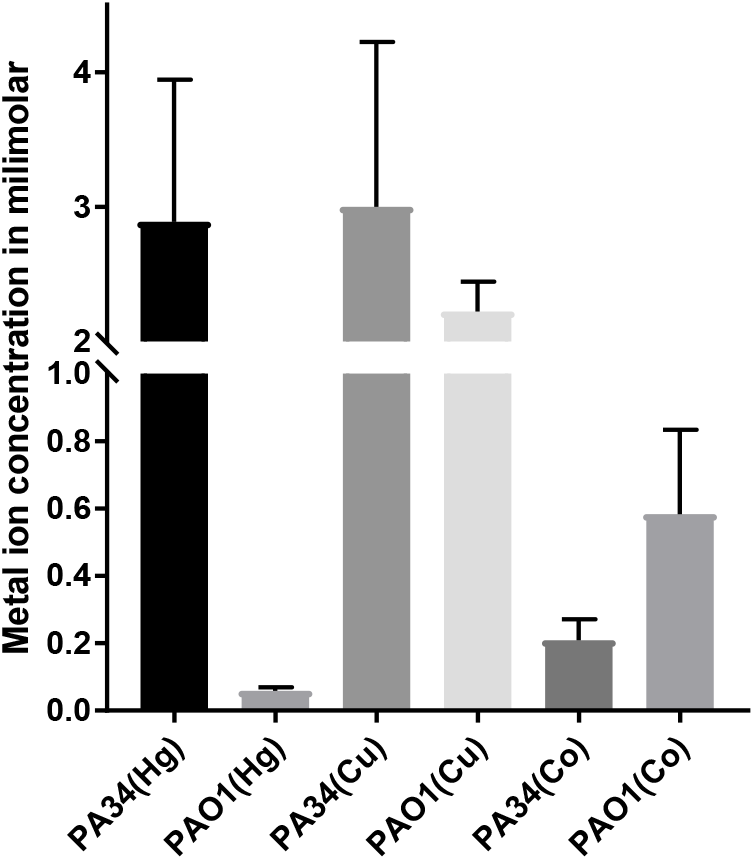
Minimum inhibitory concentration (MIC) of *P. aeruginosa* PA34 and *P. aeruginosa* PAO1 to mercury, copper and cobalt.

The *Pseudomonas* phage MP38 was identified in the new2 (flanking loci homologous to 4856/4857 of PAO1) (Fig 4B). MP38 is D3112-like phage which is a transposable and has been isolated from many clinical isolates of *P. aeruginosa* (56). We examined the presence of MP38 in genomes of different *P. aeruginosa* and found that a similar phage was integrated into VRFPA04 near the PA4204 homologue of PAO1. This confirms the result of another study (57) that has shown that transposable phages may have variable integration sites, indicating that the phage MP38 does not have any specific integration site in the genome of *P. aeruginosa*. Therefore, the new2 integration site may not be a constant RGP for *P. aeruginosa*.

Four RGPs (RGP31, RGP73, RGP9 and RGP60) of PA34 were associated with replacement of pathogenic genes and were related to lipo-polysaccharide O-antigen, pyoverdine (*pvdE*), flagella (*fliC*) and pilus (*pilA*) synthesis. Despite the replacement islands contains the same genes and being integrated into the same loci in the core genome, they highly diverse between strains (58). These components are important for interaction of a bacterium with external environments including other species or hosts and hence are under continuous selection pressure (58-61). The *pvdE* and *fliC* orthologs have been shown to vary greatly between different strains of *P. aeruginosa* (62). The replacement islands may contribute to higher virulence in PA34 although this needs to be tested.

The largest GI (RGP23) observed in PA34 was 125.1 Kbp and carried resistance genes for aminoglycosides (*AAC(3)-IId*) and tunicamycin. An insertion sequence of the family IS*1182* was observed in this GI, which may be responsible for carriage of these resistance genes. A BLASTn search revealed that the antibiotic resistance genes were best matched with those of *Acinetobacter* sp. WCHA45 plasmid pNDM1_010045 (GenBank accession number NZ_CP028560.1) and were similar to many from other members of the *Enterobacteriaceae* where IS*1182* is also present. In addition, a phage tail protein gp37, which was first identified in *Enterobacter* phage T4 (63) was found inserted into this island. This suggests that this resistance islands may be derived from phages.

## 3. Plasmid features

The Illumina^®^ and ONT^®^ reads of the whole genome sequence of PA34 were assembled into seven contigs (> 500 bp) using hybrid strategies in SPAdes. Out of seven contigs, contigs 6 and 7 had a G+C content that less than that of other contigs and showed a significant match with plasmids in the NCBI database. This suggests that contigs 6 and 7 are in fact plasmids carried by PA34 and were named here as pMKPA34-1 and pMKPA34-2. pMKPA34-1 is 95.4 kbp with 57% G+C content and pMKPA34-2 is 26.8 kbp with 61% G+C content. Automatic annotation followed by manual confirmation with BLASTn revealed 98 CDS in pMKPA34-1 and 33 CDS in pMKPA34-2. Out of 98 predicted genes in pMKPA34-1, 46 were predicted to encode proteins with unknown functions.

### 3.1 General features of pMKPA34-1

The putative plasmid pMKPA34-1 contains a replication gene (*repE*), a chromosome partition gene (*smc*), a plasmid stabilization gene (*parB*) and a plasmid conjugal transfer mating pair stabilization protein (*traN*). These genes may help replication, maintenance and transfer of the plasmid (64, 65) (Fig 7). Based on BLASTn searches against the NCBI plasmid database, pMKPA34-1 had the best match (31% query cover with 94% identity) with plasmid pLIB119 of *P. stutzeri* strain 19SMN4 (GenBank accession number CP007510), followed by *Citrobacter freundii* strain 18-1 plasmid pBKPC18-1 (23% query cover with 99% identity) (Fig. 7). The *parA, smc* and *repE* genes display high similarity to the corresponding genes of pLIB119. The presence of the *traN* gene, which encodes an outer membrane protein and is essential for F-mediated bacterial conjugation (65), indicates the potential exchangeability of pMKPA34. This transfer system is common amongst bacterial plasmids notably in *Enterobacteriaceae* (66) and its presence in pMKPA34-1 may indicate that the plasmid could be exchanged with members of the *Enterobacteriaceae*.

**Fig 7.**
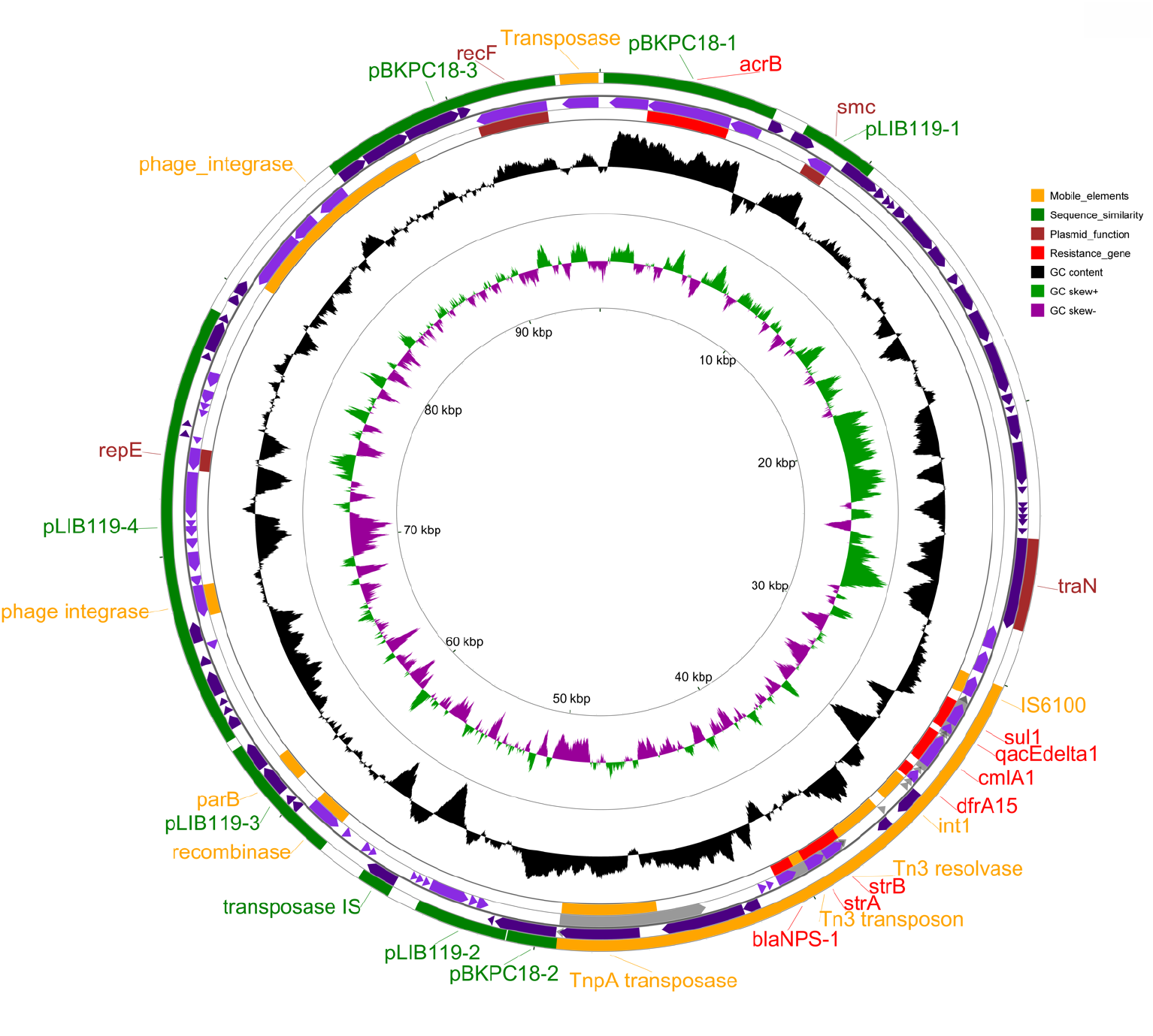
The map of *P. aeruginosa* PA34 plasmid pMKPA34-1. Circles from outer to inner represent; circle 1 and 4: positions of the matched regions representing mobile elements (orange), similarity with other plasmids (green), plasmid functions (marron), and resistance genes (red). Circle 2. Coding Sequence (CDS) in plus strand (purple), Circle 3. positions of CDS in minus strand (purple), Circle 4. G+C content and deviation from the average, Circle 5. G+C skew in green (+) and purple (–) and Circle 6. scale in kbp.

Transposase, integrase and recombinase genes formed a major portion of the mobile genetic elements in pMKPA34-1. ISsage analysis identified two putative prophage integrases, and site-specific recombinases (*xerC* and *xerD*) of the tyrosine family. The XerC-XerD complex helps segregate chromosomes at cell division and contributes to the stability of plasmids (67). It should be noted that this complex is not present in pLIB119 (Fig 7).

Plasmid pMKPA34-1 carries at least six antibiotic resistance genes. Five resistance genes (trimethoprim; *dfrA15*, chloramphenicol; *cmlA1*, aminoglycosides; *APH(3’’)-Ib/APH(6)-Id*, and beta-lactam; *bla* _NPS-1_) are located in the Tn*3*-like transposon which also carries a class I integron (In1427) possessing *dfrA15* and *cmlA1* (13). Additionally, pMKPA34-1 carries a multi-drug efflux gene (*acrB*). The *acrB* gene has high similarity to the corresponding gene of pBKPC18-1 (GenBank accession number CP022275), a resistance plasmid isolated from *Citrobacter freundii* strain 18-1. This efflux pump is similar to *acrA* and *acrB* of *E. coli* and responsible for resistance to hydrophilic compounds that include disinfectants (68). The results suggest these movable resistance genes may be related to enteric bacteria.

### 3.2 General features of pMKPA34-2

The plasmid pMKPA34-2 lacks identifiable replication genes. However, the genome coverage statistics (109x vs. 50x) (Table 1) suggests it may have a higher copy number than the pMKPA34-1. pMKPA34-2 may use alternative mechanisms for replication. In the BLASTn search against the NCBI plasmid database, pMKPA34-2 was best matched (29% query cover with 88% identity) with plasmid pSSE-ATCC-43845 of *Salmonella enterica* subsp. *enterica* serovar Senftenberg strain 775WP (GenBank accession number CP016838), followed by plasmid tig00000727 of *Klebsiella pneumoniae* strain AR_0158 (GenBank accession number CP021699) (14% query cover with 99% identity) (Fig 8). Moreover, pMKPA34-2 carried the series of genes (*tnsA, tnsB, tnsC, tnsD* and *tnsE*) that are associated with transposition and are related to Tn7 transposons, a phage integrase and a putative resolvase. Upstream of these mobile genetic elements, a multi drug export protein gene (*mepA*) was identified. MepA is multi drug efflux transporter of MATE (Multidrug And Toxic Compound Extrusion) family whose role in *P. aeruginosa* is not well understood.(69). However, MepA has been associated with resistance to many antibiotics and microbicidal dyes (crystal violet and ethidium bromide) in *Staphylococcus aureus* (70). Indeed, the presence of the *mepA* gene in this plasmid and its association with mobile genetic elements suggest that this gene may have been acquired through horizontal gene transfer.

**Fig 8.**
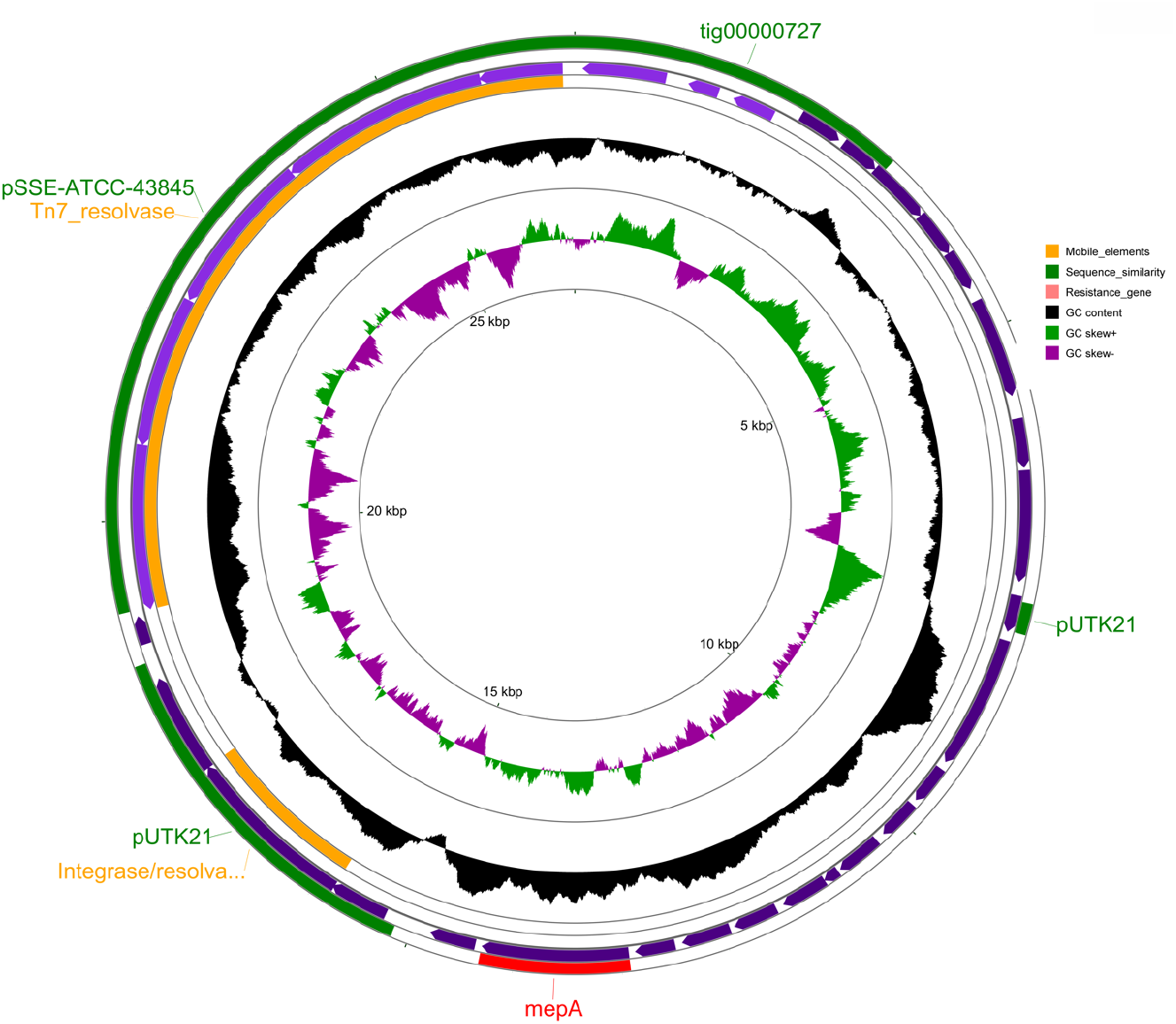
The map of *P. aeruginosa* PA34 plasmid pMKPA34-2. Circles from outer to inner represent; circle 1 and 4: positions of the matched regions representing mobile elements (orange), similarity with other plasmids (green) and resistance genes (red). Circle 2. CDS in plus strand(purple), Circle 3. positions of CDS in minus strand (purple), Circle 4. G+C content and deviation from the average, Circle 5. G+C skew in green (+) and purple (–) and Circle 6. scale in kbp.

## Conclusions

The large accessory genome of *P. aeruginosa* strain PA34 indicates that this strain has a diverse genomic structure. The strain harbours twenty-four genomic islands and two plasmids that carry various metal and antibiotic resistance genes as well as several genes associated with virulence. This may be associated with the observance of higher antibiotic resistance, mercury tolerance and in-vitro cytotoxicity of PA34. The *in-silico* analysis showed that six antibiotic resistance genes were present in two different plasmids and one antibiotic resistance gene plus various mercury resistance genes were integrated into different GIs and these resistance genes have sequence similarities with that of either other environmental or enteric bacteria. Furthermore, the genome of the PA34 has been integrated by phage element (gp37) that has its origin in enteric bacteria and carried aminoglycoside resistance gene (*AAC(3)-IId*). These findings suggest that resistance and virulence in PA34 may have evolved due to environmental selection pressure where organisms acquire traits to survive predation by other inhabitants. Thus acquired traits enhance the pathogenesis process in human. Given the eye is susceptible to being infected by environmental strains of *P. aeruginosa*, examination of a larger number of eye isolates may be necessary to uncover any additional acquired genetic features associated with microbial keratitis. This will help our understanding of different aspects of *Pseudomonas* keratitis.

## Acknowledgements

The authors would like to acknowledge Associate Professor Federico Lauro of the Asian School of the Environment and the Singapore Centre for Environmental Life Sciences Engineering (SCELSE) for providing Oxford Nanopore Technology facility for DNA sequencing. We are also thankful to UNSW high performance computing facility KATANA for providing us cluster time for data analysis.

## Author Contributions

D.S. designed the study, performed the experiments, analysed the data and wrote the drafts of the article. A.K.V. supervised D.S. and edited the article. G.S.K. helped with MinION sequencing. S.A.R. contributed to the design and implementation of the research, to the analysis of the results and edited the article. M.W. devised the project, developed the theoretical framework, edited the article.

